# Covering the bases: population genomic structure of *Lemna minor* and the cryptic species *L. japonica* in Switzerland

**DOI:** 10.1101/2024.02.14.580260

**Authors:** Marc W. Schmid, Aboubakr Moradi, Deborah M. Leigh, Meredith C. Schuman, Sofia J. van Moorsel

## Abstract

Duckweeds, including the common duckweed *Lemna minor*, are increasingly used to test eco-evolutionary theories. Yet, despite its popularity and near-global distribution, the understanding of its population structure (and genetic variation therein) is still limited. It is essential that this is resolved, because of the impact genetic diversity has on experimental responses and scientific understanding.

Through whole-genome sequencing, we assessed the genetic diversity and population genomic structure of 23 natural *Lemna* spp. populations from their natural range in Switzerland. We used two distinct analytical approaches, a reference-free kmer approach and the classical reference-based one. Two genetic clusters were identified across the described species distribution of *L. minor*, surprisingly corresponding to species-level divisions. The first cluster contained the targeted *L. minor* individuals and the second contained individuals from a cryptic species: *Lemna japonica*. Within the *L. minor* cluster, we identified a well-defined population structure with little intra-population genetic diversity (i.e. within ponds) but high inter-population diversity (i.e. between ponds). In *L. japonica*, the population structure was significantly weaker and genetic variation between a subset of populations was as low as within populations.

This study revealed that *Lemna japonica* is more widespread than previously thought. Our findings signify that thorough genotype-to-phenotype analyses are needed in duckweed experimental ecology and evolution.

## Introduction

Intraspecific genetic diversity is a key indicator for any species’ potential to adapt under global change because it provides the raw material for adaptation (Hughes & Stachowicz, 2004; Des Roches *et al*., 2018). Intrapopulation genetic diversity arises from novel mutations and, in plants, predominantly through sexual reproduction. Yet, It is known that many clonally reproducing plant species retain genetic diversity, i.e., plant populations are often multiclonal and clones are highly population-specific (Ellstrand & Roose, 1987). But how this impacts eco-evolutionary findings remains largely unclear, as molecular markers sensitive enough to characterize finer-scale variation within clonal species and populations have only recently become accessible (Ehrlich, 1988; Anisimova, 2019). Thus, practical examples of intraspecific genetic diversity impacts lag behind our theoretical understanding of its importance. Studies to fill this gap are crucial, especially when they provide a foundation to elucidate the adaptive significance of genetic diversity in changing environments.

Researchers increasingly rely on model systems like duckweed to gain insight into species responses to environmental change. Duckweeds (*Lemnoideae*) are a group of flowering aquatic plants in the Araceae family that float on the surface of still or slow-moving freshwater bodies (Landolt, 1986). They are monocotyledonous, represented by 37 species (Sree *et al*., 2016), including an interspecific hybrid, *Lemna japonica*, with *Lemna turionifera* and *Lemna minor* as parents (Braglia *et al*., 2021b). *Lemna* spp. often occur at high densities in ponds, streams and ditches in a wide range of environmental conditions (Landolt, 1975), including polluted waters. They disperse through wind, water, birds, and, increasingly, humans’ activities. Duckweeds are of high interest for applied conservation in wastewater treatment, so-called phytoremediation (Sekomo *et al*., 2012). They also hold commercial potential as a protein source for food products (e.g., as a soy substitute, Sree *et al*., 2016) and as biofuel. *Lemna* spp. have the potential to be used for revealing plant defense strategies, genome maintenance and other complex plant biochemical processes (Acosta *et al*., 2021). Duckweeds, such as *L. minor*, also have a long history of use as model organisms in fundamental research and ecological experiments due to their small size, clonal propagation and fast clonal generation time (as little as three days) (Clatworthy & Harper, 1962; Hart *et al*., 2019). Recently they have increased in popularity as model system for experimental evolution, as well as evolution of species interactions (Laird & Barks, 2018; Hart *et al*., 2019; O’Brien *et al*., 2020b,a; Tan *et al*., 2021; Lanthemann & van Moorsel, 2022).

*L. minor* reproduces vegetatively by budding, with occasional sexual reproduction through flowering described in wild populations (Hicks, 1932; Landolt, 1986). The actual frequency of flowering is unclear and likely underestimated, because the very small flower is hardly visible by eye (Ziegler *et al*., 2023). Pollination of flowers occurs through wind, water, or small animals or by direct flower contact. It is still unknown whether *L. minor* is self-compatible, but self-pollination has been shown to result in sterility in the closely related species *L. gibba* (Fu *et al*., 2017). Despite the dominance of asexual reproduction, *L. minor* may thus maintain relatively high levels of intraspecific genetic diversity for a predominately clonal species (Vasseur *et al*., 1993; Ho, 2018; but see Jordan *et al*., 1996). Importantly, there is evidence suggesting standing intraspecific genetic variation impacts fitness within populations (Jewell & Bell, 2023). In allozyme loci, intraspecific genetic diversity of duckweed is low within populations (i.e. individual water bodies like ponds) but between populations can be highly differentiated (Cole & Voskuil, 1996). Populations in close geographic proximity can harbour very different genotypes or clones (Vasseur *et al*., 1993; Ho, 2018; Hart *et al*., 2019; Tan *et al*., 2021) and may even be locally adapted to different environmental conditions (e.g. copper pollution, Roubeau Dumont *et al*., 2019). Recent studies have begun to leverage the substantial phenotypic variation in *L. minor* (Hart *et al*., 2019; Roubeau Dumont *et al*., 2019; Tan *et al*., 2021) but the intraspecific genetic variation, i.e. presence of different alleles or the population genetic structure which may drive such phenotypic variation, is still not fully understood due to a lack of high resolution population genetic studies on the species.

Importantly, it is widely assumed that while *L. minor* populations are genetically differentiated between sites they are not differentiated within site (Cole & Voskuil, 1996). However, recent evidence found that multiple clones can co-occur in a single pond (Bog *et al*., 2022). Discrepancies in previous findings may be due to the fact that many genetic studies have examined a limited portion of the species’ range, and importantly, used low-resolution genetic markers, which are not intercomparable, such as allozyme loci (Vasseur *et al*., 1993; Cole & Voskuil, 1996), ISSR markers (El-Kholy *et al*., 2015), Amplified Fragment Length Polymorphisms (Bog *et al*., 2022) or microsatellite loci (Hart *et al*., 2019; Kerstetter *et al*., 2023; Usui & Angert, 2024). Only recently genomic tools have been applied to this model system, a genotyping-by-sequencing study identified three genetically distinct populations of *L. minor* within a circa 160 km-transect in Canada (Senevirathna *et al*., 2023).

Whole-genome sequencing (WGS) is a very powerful tool, capable of revealing complex or fine population structures in species with varying reproductive systems that includes geographic clonality (Stauber *et al*., 2021) or weak population structure (e.g., Kersten *et al*., 2021). In this study, we used WGS to quantify the intra-specific genetic diversity of *L. minor* present in the species’ described distribution Switzerland to understand the structure and patterns in its genetic diversity and gain insights into how this could impact eco-evolutionary research. This is particularly pertinent because the mutation rate for the related duckweed species *Spirodela polyrhiza* is low (Wang *et al*., 2014; Ho *et al*., 2019). Though mutation rates remain unknown for *L. minor*, they are thought to be equally low and thus evolution in *L. minor* during experiments will arise from standing genetic variation. Knowing whether this standing genetic variation can be manipulated by using different locally sourced ecotypes (e.g., as done in Hart *et al*., 2019; van Moorsel, 2022) or by collecting a large number of individuals from one site, is critical information for duckweed researchers. This led us to specifically test the hypothesis that intraspecific genetic diversity within a site is low or virtually absent, but large between sites.

Many species of the duckweed family are so morphologically similar they are virtually impossible to distinguish with morphology alone (De Lange & Pieterse, 1973; Landolt, 1975). Consequently, there are also doubts about whether the species has been correctly identified in past experimental studies. A recent screening of 100 clones from the Landolt duckweed collection found that several populations of *L. minor* had been misidentified based on morphological criteria (Braglia *et al*., 2021a). A study in Canada has revealed the presence of a cryptic species (*L. turionifera*) using DNA barcoding with species-specific primers (Senevirathna *et al*., 2021). *Lemna minor* intraspecific diversity may have thus previously been overestimated due to the inclusion of cryptic species (Jordan *et al*., 1996). Within Switzerland, range maps indicate a prevalence of *L. minor* in Switzerland but very few records of closely related *Lemna* species (e.g., morphologically similar *Lemna japonica*, *L. minuta*, *L. gibba* and *L. turionifera*). Difficulties surrounding correct species identification are likely leading to incorrect distribution maps, and there is a need for large-scale genetic screening of wild *Lemna* populations (Volkova *et al*., 2023). In line with this, our second research question was to find out whether there are cryptic species present in Switzerland.

## Materials and Methods

### Sample collection

*L. minor* samples were collected opportunistically from 23 sampling sites (hereafter “wild populations”) across a 200 km range within Switzerland and one location in France during July to September of 2021 and 2022 (Fig. 1, Table 1). Sites ranged from 5 km to > 100 km apart, with some separated by mountain ranges (Fig. 1). Within sites, several microsites (samples) were collected (see also Supplemental figure S1). Given that all individuals of a sample grew close to each other, we expected that each sample contained only single clones.

**Figure 1.**
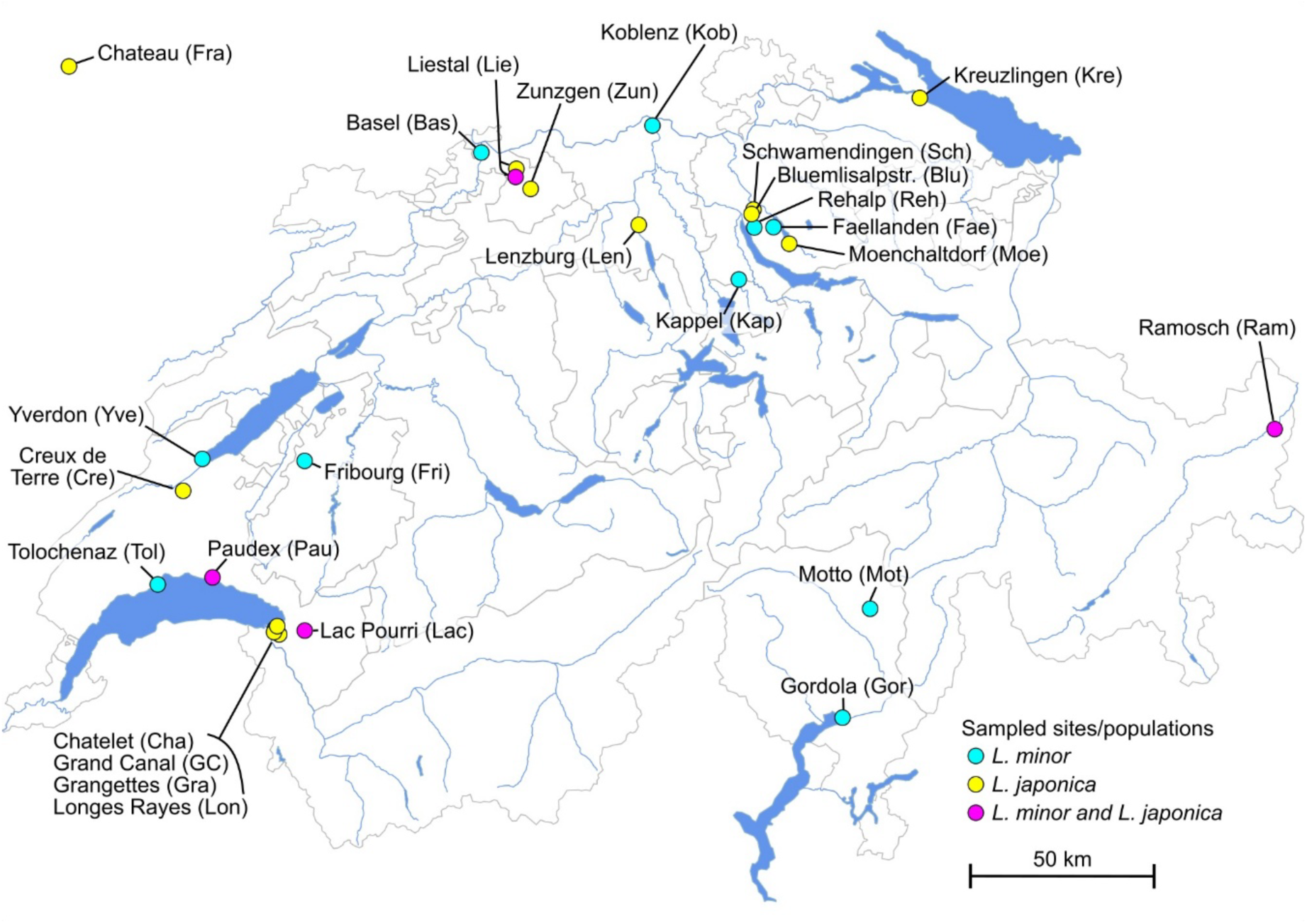
Sampling sites. Cyan for sites with only *L. minor* individuals, yellow for sites with only *L. japonica* individuals, magenta for sites with individuals from both species. Species assignment based on the kmer-approach.

**Table 1.** Sampled sites, collection dates and number of samples included in the data analyses.

| Site | Location (canton) | Lat | Lon | Date collected | N |
| --- | --- | --- | --- | --- | --- |
| Kob | Koblentz, AG | 47.60092 | 8.225556 | 07.08.2021 | 3 |
| Yve | Yverdon, VD | 46.79745 | 6.633209 | 07.08.2021 | 5 |
| Mot | Motto, TI | 46.42877 | 8.967596 | 08.08.2021 | 6 |
| Ram | Ramosch, GR | 46.83374 | 10.40119 | 08.08.2021 | 3 |
| Reh | Rehalp, ZH | 47.3524 | 8.583125 | 18.08.2021 | 3 |
| Fae | Fällanden, ZH | 47.3524 | 8.651295 | 19.08.2021 | 2 |
| Kap | Kappel, ZH | 47.2276 | 8.525658 | 26.08.2021 | 2 |
| Moe | Mönchaltorf, ZH | 47.3118 | 8.706101 | 09.09.2021 | 5 |
| Zun | Zunzgen, BL | 47.45058 | 7.790417 | 07.09.2021 | 3 |
| Lie | Liestal, BL | 47.49902 | 7.739983 | 08.09.2021 | 6 |
| Bas | Basel, BL | 47.53847 | 7.61535 | 09.09.2021 | 3 |
| Len | Lenzburg, AG | 47.36212 | 8.173317 | 17.09.2021 | 1 |
| Fri | Fribourg, FR | 46.7946 | 6.992633 | 19.09.2021 | 1 |
| Lon | Longes Rayes, VD | 46.38983 | 6.892783 | 15.10.2021 | 3 |
| Cha | Châtelet, VD | 46.37607 | 6.907367 | 03.10.2021 | 5 |
| GC | Grand Canal, VD | 46.3814 | 6.889267 | 03.10.2021 | 1 |
| Gra | Grangettes, VD | 46.39573 | 6.900383 | 15.10.2021 | 3 |
| Tol | Tolochenaz, VD | 46.49357 | 6.482117 | 14.10.2021 | 1 |
| Cre | Creux de Terre, VD | 46.71997 | 6.567183 | 14.10.2021 | 5 |
| Gor | Gordola, TI | 46.1676 | 8.864586 | 27.10.2021 | 5 |
| Fra | Chateau, France | 49.06114 | 1.738231 | 03.07.2022 | 1 |
| Blu | Blümlisalpstrasse, ZH | 7.388018 | 8.547354 | 19.09.2022 | 2 |
| Kre | Kreuzlingen, TG | 47.65712 | 9.180429 | 19.09.2022 | 1 |
| Sch | Schwamendingen, ZH | 47.39552 | 8.581181 | 21.09.2022 | 6 |
| **Landolt collection samples** |  |  |  |  |  |
| Strain No. | Species | Source location | Lat, Lon | Year collected | N |
| 9967 | *Lemna minor* | Bätzimatt, SZ | 47.21544,<br>8.941524 | 2019 | 1 |
| 9978 | *Lemna japonica* * | Oberegg, AI | 47.41857,<br>9.549893 | 2014 | 1 |
| 9969 | *Lemna japonica* * | Heiden, AR | 47.436389,<br>9.556944 | 2014 | 1 |
| 9983 | *Lemna japonica* | Meilen, ZH | 47.287669,<br>8.680481 | 2020 | 1 |
| 9965 | *Lemna japonica* * | St. Gallen, SG | 47.471269,<br>9.530747 | 2019 | 1 |
| 9700 | *Lemna minuta* | Rotterdam, NL | NA | na | 1 |
| 9478 | *Lemna turionifera* | Raciborz, POL | NA | na | 1 |

Note: \*species annotation sensu Braglia et al. (2021).

For each sample, 80 plant individuals were flash-frozen in liquid nitrogen. In addition to the wild populations, several *Lemna* clones from the Landolt duckweed collection were sampled and sequenced (also 80 individuals per sample). Each strain represented true clones with a recorded history in lab culture and scientific use. To act as reference outgroups, we included the following strains from candidate cryptic species: *Lemna minuta* (strain 9700 collected in Rotterdam, the Netherlands), *L. japonica* (strain 9983 collected in Meilen, Switzerland) and *L. turionifera* (strain 9478 collected in Raciborz, Poland). We aimed to include *L. gibba*, however, the strain we used (9965 collected in St. Gallen, Switzerland) was re-classified during our study as *L. japonica* (Braglia *et al*., 2021a) based on Tubulin Based Polymorphisms (TBP). Furthermore, as described by Braglia and colleagues (2021), two of the used *L. minor* strains were reclassified as *L. japonica* (strains 9969 and 9978). All samples of the wild *L. minor* populations and the duckweed stock populations were kept at −80°C for downstream applications. In total, 87 samples (each based on 80 pooled plant individuals) that yielded high-quality data for genomics were sequenced (see Table 1).

### DNA extraction

For DNA extraction, 30 mg of the frozen tissue consisting of several fronds was weighed into 1.5 ml tubes (Eppendorf, Hamburg, Germany) with two 3 mm metal beads and ground into a fine powder with a TissueLyser II (Qiagen, Germany) at 30 Hz for 2 min. The extraction was performed using the Norgen Plant and Fungi genomic kit (Norgen Biotek, Thorold, ON, Canada) with minor modifications. Specifically, 750 μL of lysis buffer supplemented with 1 μL RNAse A (DNase-free, 100,000 units/mL in 50 % glycerol) and 3 μL Proteinase K (>600 u/ml ∼ 20 mg/ml in 10 mM Tris-HCl pH 7.5, containing calcium acetate and 50% v/v glycerol) was added to each tube and vortexed vigorously. Then, 150 μL of Binding Buffer I was added to each tube, vortexed and incubated for 5 min on ice. The rest of the extraction procedure was conducted according to the manufacturer’s protocol. Finally, the purified DNA was eluted in 100 μl of elution buffer and stored at −20°C. Quantification and quality control of the purified DNA was performed by Qubit dsDNA HS Assay Kits (Thermo Fisher Scientific) and gel electrophoresis. Library preparation and sequencing was conducted by Novogene (UK) using Illumina NovaSeq PE150 paired-end sequencing (Novogene, Cambridge, UK).

### Data processing

#### Short read processing

Short reads were first de-duplicated with NGSReadsTreatment (version 1.3, Gaia *et al*., 2019) and quality trimmed with fastp (version 0.20.0, Chen *et al*., 2018).

#### K-mer based approach

To process samples for population genetic analysis in a reference free manner, we used a recently described pipeline (Voichek & Weigel, 2020), github.com/voichek/kmersGWAS). Analyses were conducted using the default k-mer size of 31. The kinship matrix generated with the pipeline was converted to a distance matrix (1-kinship) to enable comparability across analyses. It should be noted that we did not intend to provide a full comparison between k-mer and alignment-based pipelines but to use a reference-free approach to assist with species misassignments.

#### Reference-based approach

We used the reference genomes of either *L. minor* strain 7210 or the diploid *L. japonica* strain 9421 (Ernst *et al*., 2023). The *L. minor* assembly includes 21 chromosomes (362.1 million bp, all sizes haploid), 30 unassigned scaffolds (0.9 mio bp), and the plastid genomes (0.4 mio bp). The *L. japonica* assembly includes 21 chromosomes from the *L. minor* parent (347.0 mio bp), 21 chromosomes from the *L. turionifera* parent (420.3 mio bp), 85 unassigned scaffolds (3.2 mio bp), and the plastid genomes (0.4 mio bp). Reads were aligned using bowtie2 (version 2.4.5, with a minimal mapping quality of 10, Langmead & Salzberg, 2012). Runs of samples with more than one sequencing run were merged with samtools (version 1.18, Danecek *et al*., 2021). SNPs were identified with bcftools (version 1.18, Danecek *et al*., 2021) and filtered for a minimal quality of 20. SNPs were further filtered for a coverage between 10 and 1,000 within at least 80 % of all samples. Pairwise genetic distances were calculated as the fraction of alleles that differed between two individuals, while ignoring SNPs with pairwise missing data. After identification of the *L. minor* and *L. japonica* subgroups (see below), the subgroups were divided and treated separately. Each cluster was re-analysed as described above using the species-specific genome.

#### Non-plant DNA

To assess whether the samples contained sequences from other organisms, we scanned all reads with the metagenome classifier tool CLARK (version 1.2.6.1, Ounit *et al*., 2015). We used the family-level databases of fungi, bacteria, and viruses (set_targets.sh options: bacteria viruses fungi --family) and a k-mer size of 31 (classify_metagenome.sh options -k 31 -m 2 --ldm --light). Abundances were obtained with default settings (estimate_abundance.sh). The fraction of reads that were assigned during the process was extracted from the output of classify_metagenome.sh. We filtered for families with at least 10 sequence counts in 10 samples and used DESeq2 (version 1.24.0, Love *et al*., 2014) to obtain log2(x+1) transformed normalized counts that were used to calculate pairwise Manhattan distances between the samples. Differential abundance of families between *L. minor* and *L. japonica* individuals was done as described previously (Schmid *et al*., 2019) and families with an FDR < 0.05 and a log2 fold-change of at least 2 were considered to be differentially abundant.

### Data analysis and visualization

#### SNP-wise test for association with genetic clusters (“assignment markers”)

To test for associations of a genetic variant with a group of samples (i.e., the two genetic clusters which corresponded to the two species *L. minor* and *L. japonica*), we ran an ANOVA for each individual SNP using R (version 4.2.3, i.e., we ran a GWAS using a simple GLM). The genetic variants in case of bi-allelic SNPs have three possible levels corresponding to “homozygous reference”, “heterozygous”, and “homozygous alternative”: 0/0, 0/1, and 1/1. Besides this three-level factor, we also used two two-level factors for comparisons of one homozygous state to the other two states (factor A: 0/0 *vs.* 0/1 and 1/1; factor B: 1/1 *vs.* 0/1 and 0/0). We then only considered the factor with the best fit. Factorial data was tested with ANOVA(glm (formula, family = quasibinomial(“logit”)), test = “F”)) and the effect size was calculated as the percent of deviance (%-DEV) explained by the variant. *P*-values were adjusted for multiple testing to reflect false discovery rates (FDRs). Associations with an FDR below 0.05 and %-DEV of at least 80% were considered significant. We chose the high %- DEV cut-off to only include cases that were clearly and consistently different between the two species. Given the relatively large sample size, only filtering by FDR would also yield SNPs that are specific to for example one population and not the “entire” species.

#### Differences in variation among populations

We tested whether the intra-population variation differed among populations using the function betadisper() from the R-package vegan (version 2.5.7, Dixon, 2003). The function was also used to extract the distances to the overall species’ centroid by treating the entire species as one population (used for plotting).

#### Isolation by distance

Isolation by distance, which involved the correlation between genetic distance matrices and the correlation of genetic distances with physical distances, was tested with Mantel tests using 9,999 permutations with the R-package vegan (version 2.5.7, Dixon, 2003). Correlations between genetic and physical distances were also tested with an ANOVA (anova(lm()). In both cases, physical distances in meters were log10 transformed.

#### Visualizations

Heatmaps were generated using the distance matrices and the function gl.plot.heatmap() from the R-package “dartR” (version 1.0.2, Gruber *et al*., 2018). The genetic composition of the individual samples in Figure 3A was inferred using fastSTRUCTURE (version 1.0, Raj *et al*., 2014) as described in the online manual (rajanil.github.io/fastStructure). The number of populations was set to K = 8 (*L. minor*) and K = 4 (*L. japonica*) because this was the number of expected populations based on the distance visualizations (Figure 3). Supplemental figure S4B was done using the function heatmap.2 from the R-package gplots (version 3.1.1, Warnes *et al*., 2024). All other plots were done in R using base graphics.

## Results

### Three different approaches recovered two species

The kmer approach identified 1,357,142,099 variants in the kmer table. Alignment to the *L. minor* reference genome (strain 7210) and SNP calling with bcftools (Danecek *et al*., 2021) yielded 10,910,630 SNP’s in total, out of which 1,343,857 SNPs had a coverage of at least 10 in at least 80 % of all individuals. Alignment to the *L. japonica* reference genome (strain 9421) and SNP calling yielded 10,067,012 SNPs, out of which 1,511,714 SNPs passed the coverage filter. We created kinship matrices based on all three approaches (Fig. 2A) and recovered two distinct genetic clusters. The included *L. minor* sample from the Landolt duckweed collection with known species attribution (based on Tubulin polymorphism, Braglia et al 2021) clustered together with one subgroup, whereas the collections’ *L. japonica* samples clustered with the other subgroup (see grouping in Fig. 2A). To test whether these two clusters could be attributed to two species, we used three additional approaches: 1) alignment rate comparison, 2) identification of cluster-specific SNPs, and 3) comparison of alternative allele depths using also publicly available data (Ernst *et al*., 2023).

**Figure 2.**
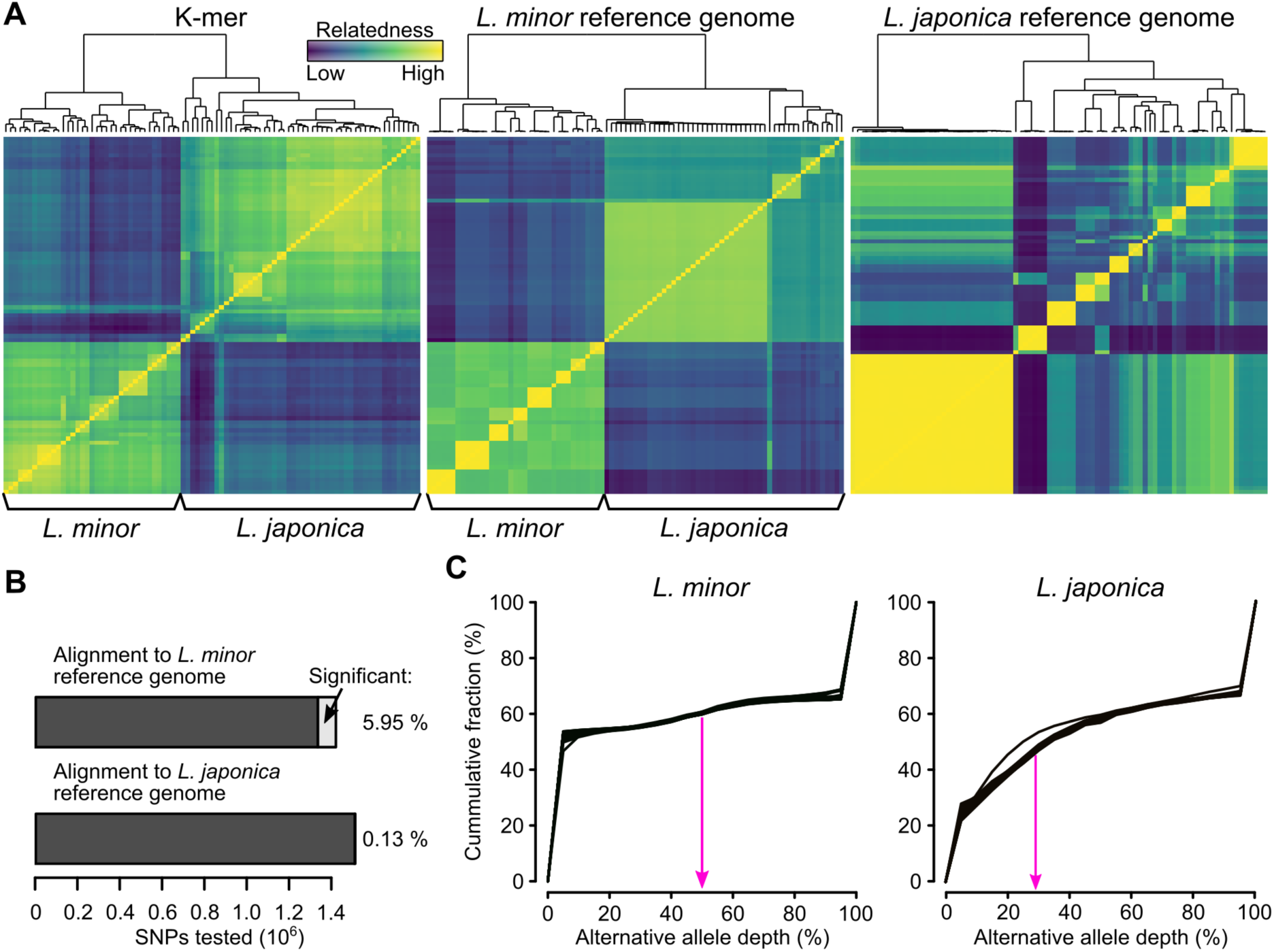
Analysis with all samples. A) Kinship matrices based on kmers, alignment with *L. minor* reference genome (Lm7210), and alignment with *L. japonica* reference genome (Lj9410). Species were assigned based on the kmer-approach and match to the approach with *L. minor* as reference genome. B) Test for genetic differentiation between *L. minor* and *L. japonica* on the level of individual SNPs. Significance level was set to FDR < 0.05 and %-deviance explained by the species > 80 %. Most differences were homozygous *versus* heterozygous differences: 99.73 % and 80.69 % in case of the *L. minor* and the *L. japonica* reference genome, respectively. C) Allele frequencies in *L. minor versus L. japonica* individuals using the *L. minor* reference genome. *Lemna minor* individuals have an enrichment of non-homozygous frequencies at 50 %. In contrast, *L. japonica* individuals have an enrichment of non-homozygous frequencies around 33 % (see also Supplemental figure S3 with data from the publicly available reference genomes).

#### Differences in alignment rates

Alignment rates for the *L. japonica* reference genome ranged from 50 – 90 %, except for the *L. minuta* sample which had an alignment rate of 2.9 %). Rates for the *L. minor* genome were generally lower for all samples, but there was a marked difference among samples (Table S1, Supplemental figure S2). While the alignment rates of the *L. minor* samples were on average only 10.2 % lower (range: 5.6 – 13.9 %), alignment rates of *L. japonica* samples were about 26.1 % lower (range: 16.3 - 32.4 %). As a comparison, the alignment rate of the *L. turionifera* sample, whose genome is only represented in the *L. japonica* reference, was 66.1 % lower.

#### Species assignment markers

We tested all SNPs that passed the filter for significant association with either of the clusters (i.e. species) using generalized linear models (GLMs) followed by multiple testing corrections and stringent filtering (FDR < 0.05 and %-deviance explained > 80 %). While using the *L. minor* reference, 5.95 % of all SNPs were highly differentiated between the two clusters. In contrast, with the *L. japonica* reference only 0.13 % of all SNPs showed significant differentiation (Fig. 2B). Meeting with theoretical expectations for allele frequencies between species, most differentiated markers were homozygous fixed differences (99.76 % *L. minor* reference, and 82.30 % *L. japonica* reference). The difference in the fraction of significant SNPs was probably due to the fact that parts of the genome from *L. japonica* (i.e., the copy from the parent that is not *L. minor,* hence *L. turionifera*) wrongly aligned to the *L. minor* reference sequence, and that this was corrected when using the *L. japonica* reference genome. The limited difference between the two clusters with the *L. japonica* reference genome further suggested that the copy of the *L. minor* parent in the *L. japonica* individuals was similar to the genome of the *L. minor* individuals.

#### Alternative allele ratio suggests that L. japonica is triploid

To verify the difference between the two clusters and assess the genetic composition of the *L. japonica* individuals, we extracted the “alternative allele ratio” at each SNP using the *L. minor* reference genome data (Fig. 2C). For example, one SNP might be covered with 30 reads that suggest that the base is “A” like in the reference genome, whereas another 30 reads suggest an alternative base “T”. Hence, the ratio would be 0.5 and indicate that the individual is heterozygous at this position. However, if there were only 15 reads suggesting the base “T”, the ratio would be 1:2 and indicate that the individual might be triploid with one copy being different (or hexaploid with two copies being different). Doing this for all SNPs, we indeed found an enrichment of the alternative allele ratio of 50 % in all *L. minor* individuals and an enrichment of an alternative allele ratio around 33 % in all *L. japonica* individuals (Fig. 2C). To verify this approach, we added data from the previously published *Lemna* reference genomes (Ernst *et al*., 2023), Supplemental Figure S3). Among them were three different *L. japonica* hybrids with different mixtures of the parental genomes M (*L. minor*) and T (*L. turionifera*): 1) a diploid M/T, 2) a triploid M/M/T, and 3) a triploid M/T/T hybrid. These three hybrids matched the order of the expected alternative allele ratios with the lowest in M/M/T, intermediary in M/T, and highest in M/T/T. While the expected ratios would be in theory 33, 50, and 66%, the observed ratios were slightly lower (see Supplemental figure S3). It is unclear why this was the case. Nonetheless, all *L. japonica* in our study were closest to the triploid M/M/T hybrid, indicating that *L. japonica* in Switzerland also comprises triploid M/M/T hybrids. Additional support for this hypothesis came from the average coverage of the two sub-genomes, *L. minor* and *L. turionifera*, in the *L. japonica* reference genome. While the *L*. *minor* samples had practically zero coverage on the *L. turionifera* sub-genome, the *L. japonica* samples exhibited approximately a 2:1 ratio in genome coverage between the *L. minor* and the *L. turionifera* sub-genomes (Supplemental figure 3C).

Taken together, our analyses suggest the widespread presence of a triploid *L. japonica* hybrid with two copies of the *L. minor* genomes and one copy of the *L. turionifera* genome in Switzerland. We will, therefore, refer to the two genetic clusters as *L. minor* and *L. japonica* groups.

### Finer population structure depended on the used approach

The population structure depicted in Figure 2 revealed general agreement among the three datasets. However, we highlight a few differences to demonstrate the need for caution when it is uncertain whether all individuals in a study belong to the same species. While correlations between pairwise distances were clearly significant (P_Mantel_ = 0.0001 with 9,999 permutations), they were still markedly distinct. The correlation between the kmer and the *L. minor* reference genome data sets was relatively high (Pearson correlation coefficient = 0.9). However, the correlation between *L. japonica* reference and the *L. minor* reference was lower (Pearson correlation coefficient = 0.71). The *L. japonica* reference and the kmer dataset correlated only weakly (Pearson correlation coefficient = 0.56). This discrepancy was also visible in the inconsistent placement of some *L. japonica* individuals in the three dendrograms (Fig. 2A). We ultimately used the assignment from the kmer and alignment to *L. minor* reference genome approaches, which we could verify as described above.

Furthermore, the distance values (see dendrogram and color-gradient in Fig. 2A), suggest that contrasting conclusions could be drawn. The kmer data set suggests considerable intrapopulation and interpopulation variation. The *L. minor* reference genome dataset indicates practically no intrapopulation variation but significant interpopulation genetic variation for the *L. minor* samples. Meanwhile, for the *L. japonica* group it would suggest intermediate variation within and between populations (ignoring for now the large group of similar *L. japonica* individuals which comprises multiple populations). Again, in contrast, the *L. japonica* reference genome dataset would suggest very little intrapopulation variation and large interpopulation variation for both species. The different output resulting from the kmer approach might have been partly due to the presence of environmental DNA, as we did not remove the reads assigned to eDNA prior to the kmer analysis. The difference between the two reference genomes was solely due to the presence of the *L. turionifera* sub-genome in the *L. japonica* assembly. Given that half of the samples were *L. minor* and we filtered SNPs that were not covered by at least 80 % of all samples, this filter masked much variation in the *L. japonica* group on the *L. turionifera* sub-genome while the presence of the *L. turionifera* sub-genome during the alignment lead to fewer errors in the *L. minor* sub-genome. While conclusions would not be much different for the *L. minor* samples, they would be distinct for the *L. japonica* samples. Hence, to assess the population structure of the two species individually, we split the data into two sets: 1) *L. minor* samples using the *L. minor* reference and 2) *L. japonica* samples using the *L. japonica* reference.

### Population genomic structure differed between *L. minor* and *L. japonica*

To assess the wild population structure for *L. minor* and *L. japonica*, we kept the two species separate and excluded the samples from the Landolt collection. We further excluded three samples in the analyses of the within- and between-population distances (R4, GR6, and BA1) because they were genetically implausibly far apart from their population of origin. R4 grouped with samples from another population, suggesting a mislabelling during processing. The other two samples might indeed be markedly different genotypes within their population, but at least for BA1 it seems highly unlikely because this population originated from an isolated, tiny pond in a garden surrounded by large buildings. Thus, both should be verified independently.

While it is possible that with our sampling approach we pooled different genotypes, we would expect differences in the allelic ratios in such genotype mixtures and assumed this would be highly visible. However, except for one sample, allelic ratios were highly similar between samples of a given species and were also similar to the pattern observed in the strains that were cultivated and kept as single clone (Supplemental Figure S3). Thus, it seems unlikely that we pooled more than one clone per sample (except for GR6).

Kinship matrices and dendrograms (Fig. 3A) indicated that *L. minor* populations were well structured. In contrast, *L. japonica* was split into one large cluster comprising multiple populations, one group comprising two populations, and two remaining populations. Thus, several of the originally sampled *L. japonica* populations were clearly not distinct from each other. We therefore used the grouping inferred from the kinship matrices and dendrograms to compare within- to between-population distances (Fig. 3B). In both species, intra-population differences were much smaller than between-population differences (*P* < 10^-10^). Distances within groups were on average 0.0147 (*L. minor*) and 0.0126 (*L. japonica*). In contrast, distances between groups were on average 0.2779 (*L. minor*) and 0.298 (*L. japonica*). Using the original population labels we found that intra-population variations were significantly different in *L. japonica* (*P_Mantel_* = 0.0001 with 9,999 permutations) but not in *L. minor* (*P _Mantel_* = 0.1611, Fig. 3B).

**Figure 3.**
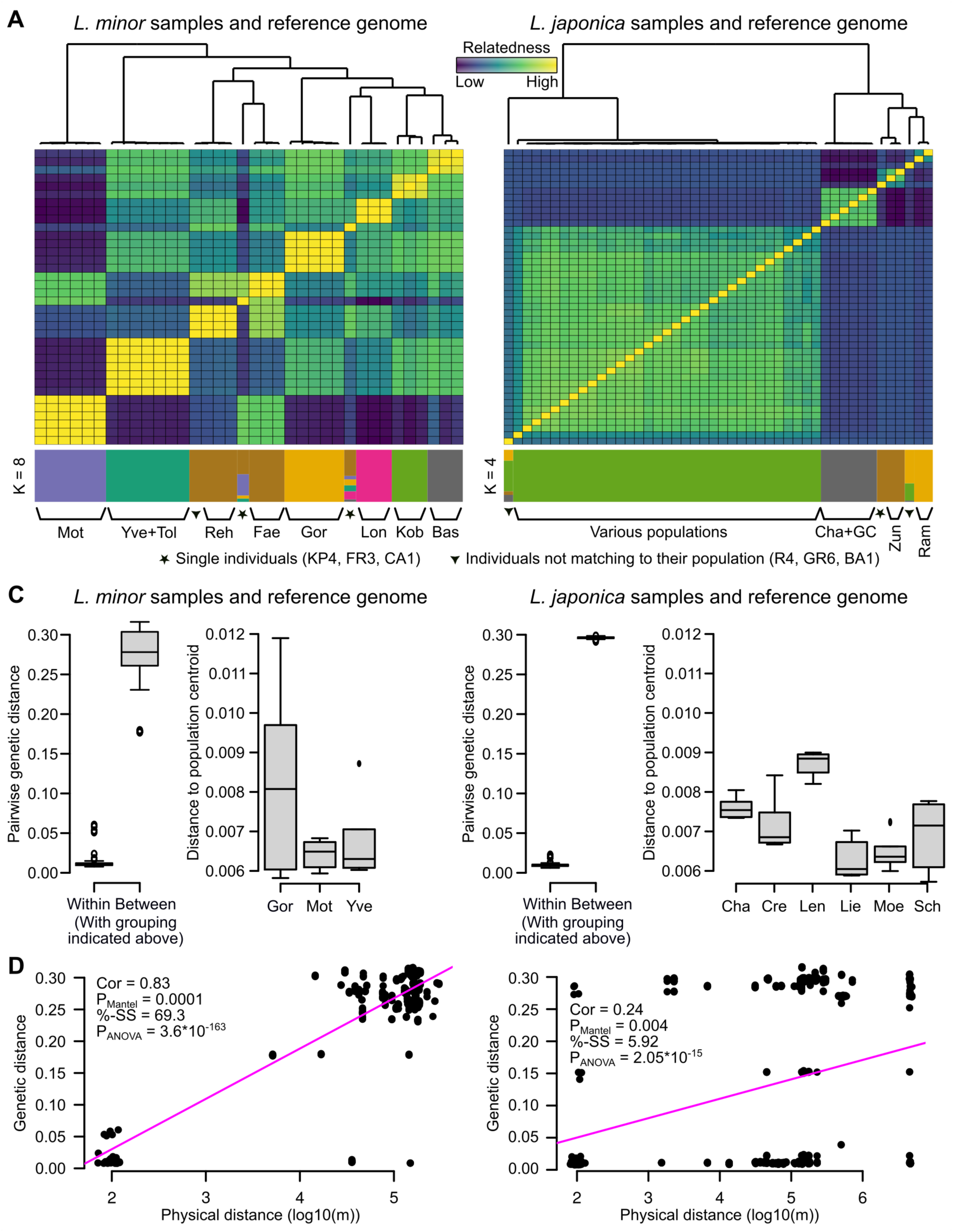
Per-species analysis. A) Kinship matrices for *L. minor* (left) and *L. japonica* (right). Whereas the populations of *L. minor* are generally well separated from each other, several populations of *L. japonica* are forming a single group. The STRUCTURE plot at the bottom visualizes the genetic composition of the individual samples. The stars below the structure plot refer to individuals that are alone in representing their population, and hence, potentially less reliable data points. The diamond refers to individuals that do not match to their population. B) Left panels with pairwise genetic distances summarized within and between groups using all groups with at least three individuals. Note that the groups were formed according to the grouping indicated in the kinship matrices above. I.e., especially in case of *L. japonica*, one group comprises multiple populations. Right panels show within population distances to the population centroid (only populations with at least four individuals are shown). C) Correlation between genetic and physical distances. The correlation was strong and highly significant in case of *L. minor*. Even though still significant in *L. japonica*, the correlation is much weaker.

We finally tested whether genetic distances correlated to physical distances using both Mantel tests and an ANOVA (Fig. 3C). The correlation was significant in both species (both *P*_Mantel_ < 0.01 and both *P*_ANOVA_ < 10^-10^) but much higher in *L. minor* (Pearson correlation of 0.83) compared to *L. japonica* (Pearson correlation of 0.24).

Considering that *L. japonica* was a hybrid with the genomes of both parents available in the assembly, we could compare the variation within these two “sub-genomes”. We, therefore, split the SNPs according to the sub-genome of origin, re-calculated the pairwise distances, and compared the distances to each other. If the two were markedly different from each other, it might indicate that the data contained samples with different parents. Due to differences in coverage (Supplemental figure S3), the number of SNPs was significantly different: 516,007 and 4,081 in the *L. minor* (M) and the *L. turionifera* (T) sub-genome, respectively. Nonetheless, pairwise distances were very similar to each other (Pearson correlation of 0.95, *P_Mantel_* = 0.0001 with 9,999 permutations). It is therefore possible that the *L. japonica* samples have a common hybrid ancestor and that the differences we observed were gained after the original hybridization. However, it is also still possible that there were multiple hybridization events but that the dissimilarity between the parents was similar.

Similar to comparing the variation between the two subgenomes, we further extracted SNPs in the mitochondrial and chloroplast genomes and re-calculated the pairwise distances using these subsets. Distances based on the plastid genomes were similar to each other (Table 2). When distances based on plastids were compared to distances based on the M and T subgenomes, correlations between the mitochondrial and the nuclear genomes were higher than correlations between the chloroplast and the nuclear genomes. However, in both cases, correlations to the M subgenome were clearly higher than those to the T subgenome (Table 2). This suggests that the female parent that contributed the plastid genomes was likely *L. minor*. This is further supported by visualizing the similarity in the plastid genomes using all samples, including publicly available data (Supplemental figure 3D). While all *L. minor* and *L. japonica* samples are similar, the two *L. turionifera* and the *L. minuta* samples are very distinct. This is especially interesting considering that the publicly available *L. japonica* samples have different subgenome compositions (MT, MMT, and MTT).

**Table 2.** Pairwise distances using only SNPs in specific sub-genomes of the *L. japonica* assembly. C: chloroplast, M: mitochondrial, M: *Lemna minor*, T: *Lemna turionifera*.

| Subgenome A | Subgenom B | Correlation | P-value |
| --- | --- | --- | --- |
| plastid_C | plastid_M | 0.929 | 0.0001 |
| plastid_C | subgenome_M | 0.471 | 0.0001 |
| plastid_C | subgenome_T | 0.297 | 0.0295 |
| plastid_M | subgenome_M | 0.742 | 0.0001 |
| plastid_M | subgenome_T | 0.578 | 0.0001 |

In summary, we found clear population structures that also reflected the physical distance between the populations, especially in *L. minor*. In *L. japonica*, the structure was less pronounced, and several populations were similar enough to be grouped into one large group exhibiting “within-population-level” genetic differences even though they were physically well separated from each other (“various populations” in Fig. 3A).

### Non-plant eDNA associated with collected samples

On average, about 6.19 % of all reads were classified as bacterial, viral, or fungal and the two species were alike (Supplemental figure S4A). In total, 555 different fungi/bacteria/virus families passed the filters, resulting in a table with varying read counts per sample and fungal/bacterial/viral family. Pairwise Pearson correlation coefficients between samples using these data were between 0.67 and 0.99. Samples were not grouped by eDNA according to populations, but to some extent by species (Supplemental figure S4B). Likewise, correlations between pairwise distances from the eDNA data to the three approaches above were limited (but *P_Mantel_* < 0.01): with the *L. minor* genome: 0.17, with the *L. japonica* genome: 0.20, and with kmers: 0.09. Thus, the eDNA in the samples did not reflect the population structure observed with the plant data. Averaged *L. minor* and *L. japonica* samples were highly similar to each other (correlation of 0.98, Supplemental figure S4C). Out of the 555 families, five showed differential abundant in *L. japonica* compared to *L. minor.* All five families were more abundant in *L. japonica* samples: *Desulfuromonadaceae*, *Fastidiosibacteraceae*, *Paludibacteraceae*, *Prolixibacteraceae*, and *Marseilleviridae*. While little is known about these families, there are differences for some families and the distances in eDNA did not simply reflect those obtained from the plant data. This suggests that a proportion of reads originated from organisms other than the studied plants. While this may be used to study the microorganisms associated with the plants, it also indicates that the kmer approach should be used carefully as it does not inherently filter against sequences coming from non-target organisms. An option might be to filter reads matching to eDNA before running the kmer analysis.

## Discussion

Within this study, we used sampling of natural sites of *Lemma minor* in Switzerland and France, coupled with whole genome sequencing, to characterise population structure and standing genetic variation of this common model system. We found expected patterns of low within site variation relative to between site differentiation in the species. Perhaps, most strikingly, we found much of the described distribution of *L. minor* actually contained *L. japonica*, with the two species also showing different patterns of differentiation.

### Higher between- than within population genetic variation in *L. minor*

*L. minor* generally displayed very low within population genetic diversity. It is noteworthy that the genetic distances reflect the fraction of alleles that differ between individuals and are calculated only on positions where at least one individual has an alternative allele. Thus, within population distances are likely overestimates. The low levels of intrapopulation differences (∼1.5 %), signal that it is likely that each site contained a clonal population. However, we could not confirm this with replication of samples due to lack of material. Differences between sites were much larger than those within sites, indicating different clones are likely present in different ponds and as a result *L. minor* shows spatial genetic structure with signals of isolation by distance. These findings are in line with previous research based on lower resolution genetic markers (Cole & Voskuil, 1996; Hart *et al*., 2019; Bog *et al*., 2022). However, more genetic variation was found here relative to the comparable geographical range reported in Senevirathna *et al*., (2023). These results indicate that when including populations from multiple sites in an experimental setting, chances are high that these combined populations contain multiple clonal genotypes. This will result in a baseline population with some standing genetic variation on which selection can act, the exact level of which will depend on the number of sites mixed and the distance between them. However, when including only individuals from one site (i.e., a single population), there is a high chance that the individuals are genetically very similar to each other, if not clonal. This difference in baseline will likely impact the obtained results (van Moorsel, 2022). Furthermore, knowledge of the standing genetic variation is needed for a mechanistic understanding of adaptive processes in experimental evolutionary studies (Barrett & Schluter, 2008; Lemmen *et al*., 2022; Usui & Angert, 2024).

### Presence of cryptic species revealed

In this study, we identified that a large number of sites were *L. japonica*. Indeed, we showed the presence of both *L. minor* and *L. japonica* is widespread in Switzerland. This was surprising, as current species range maps of *L. minor*, such as for example provided by Info Flora (https://www.infoflora.ch/de/flora/lemna-minor.html), indicate a broad distribution of *L. minor* across Switzerland but no records of *L. japonica*. Correctly attributing species within the *Lemna* family is challenging (Volkova *et al*., 2023). However, our results are supported by recent evidence, the presence of *L. japonica* in Switzerland was recently inferred based on the re-analysis of strains from the Landolt duckweed collection that showed a number of species misassignments (Braglia *et al*., 2021b). However, the large geographical extent of *L. japonica’s* distribution was unknown prior to our study. The substantial geographical range shown here indicates duckweed research should molecularly confirm the focal species. This ensures accurate conclusions are drawn and experimental findings are attributed to the correct species. Perhaps also due to erroneous species attribution, little is known on how *L. minor* and *L. japonica* vary in traits relevant for practical purposes. Recent evidence suggests that the two species may vary in functional traits, such as relative growth rate or thermal performance breath (Usui & Angert, 2024). Generally, closely related species of duckweed are known to exhibit strong differences in phenotypic traits, such as for example tolerance to pollutants (Lahive *et al*., 2011), and it remains to be tested whether this also holds true for *L. minor* and *L. japonica*.

### Population genomic structure of *L. japonica*

In contrast to *L. minor*, the population genomic structure of *L. japonica* is less clearly defined. Many sites were highly similar to each other (on the level of intrapopulation variation) and thus potentially clonal. However, this was spatially variable with some groups of sites showing levels of differentiation like the patterns seen in *L. minor*. This is in line with the hypothesis that hybridization between *L. minor* and *L. turionifera* has occurred multiple times, leading to independent lineages of *L. japonica* with different ploidy or chromosome rearrangements (Braglia *et al*., 2021a; Ernst *et al*., 2023). However, it is unlikely these have arisen recently in situ in Switzerland, because the sampling campaign did not recover any of the parent species *L. turionifera.* Furthermore, *L. turionifera* is thought to be currently and historically only present in three sites across all of Switzerland (https://www.infoflora.ch/de/flora/lemna-turionifera.html). Taken together, this suggests that the sampled *L. japonica* hybrids arose outside of Switzerland and were dispersed by metazoan activity.

We compared our *L. japonica* samples to multiple hybrid *L. japonica* lineages from three continents with different dosages of the parental *L. minor* and *L. turionifera* genomes as described in Ernst et al. (2023). It is perhaps not surprising that all the *L. japonica* isolates sampled in our study have the same MMT sub-genome configuration as the isolate from Denmark (*L. japonica* 8627). The other hybrids are from Africa and North America and have different sub-genome dosages. Broader geographical comparisons of *L. japonica* individuals with same ploidy level and configuration (M/M/T) are now needed to confirm if multiple hybridization events lead to different *L. japonica* lineages or if all the *L. japonica* individuals of the M/M/T configuration have a single common ancestor. This is relevant as different clonal lineages may behave different depending on their ancestry as has previously been observed in fungal clones of *C. parasitica* (Stauber *et al*., 2021).

Our findings suggest that all *L. japonica* samples were triploid, which may explain the weaker population structure in *L. japonica* as compared to *L. minor*. Triploidy is often associated with infertility and vegetative propagation (Ramsey & Schemske, 1998), indicating that *L. japonica* is more likely to be clonal than its diploid sister species. Polyploidization is assumed to be one of the major mechanisms of plant speciation and evolution (Herben *et al*., 2017). The relative importance may be greatest for plant species which reproduce mostly vegetatively, and thus experience less genome mixing via sexual reproduction. *L. minor* and *L. japonica* could thus present an interesting comparative system to study the evolution of sex.

It remains to be tested whether this *L. japonica* population genomic structure is ubiquitous to the species or specific to the unique geographical structure or colonisation history of Switzerland. Recent interest in this species for technical applications (Liang *et al*., 2023) certainly warrants a deeper investigation. It might also be worth to consider studying DNA methylation as this can resolve intraspecific phylogenies with a high temporal resolution (Yao *et al*., 2023).

### Conclusions

We used whole-genome sequencing to elucidate the population genomic structure of *L. minor* in Switzerland. Our analyses revealed the presence of *L. japonica* in Switzerland. Consistent with previous studies, we conclude that the morphological analysis alone may not be sufficient for accurate species attribution within *Lemna* spp. in the field. Future studies using Lemnaceae as a model system in experimental population genetics (Acosta *et al*., 2021), community ecology (Laird and Barks, 2018) and eco-evolutionary dynamics (Hart et al., 2019) or applied research should ensure the accurate identification of *L. minor*. Our study offers valuable genomic resources for researchers working on Lemna species. Whole-genome sequencing has become a cost-effective and powerful tool to describe duckweed population structure and should be used in future efforts to map species and genotype presence of Lemna species.

## SUPPORTING INFORMATION

Supporting information are available online and consist of the following:

**•** Table S1. Alignment rates and number of reads per sample.
**•** Supplemental figure S1: Overview of field sample collection procedure.
**•** Supplemental figure S2: Alignment rates for all samples using the *L. minor* and the *L. japonica* reference genome.
**•** Supplemental figure S3: Allele frequencies in *L. minor versus L. japonica* individuals using the *L. minor* reference genome and the *L. japonica* reference genome.
**•** Supplemental figure S4: Analysis of reads to bacterial, fungal, and viral sequence databases.

## ACKNOWLEDGMENTS

Many thanks to the people who assisted in the collection of the plants in the field: Sofia Vámos, Emilie Hanus, Benjamas Ramasauer, Doris Stetak, Barbara Känel Hofstetter, Gregor Kozlowksi, Daniel Küry and Bernhard Schmid. Thanks to Walter Lämmler from the Landolt collection for providing the *Lemna* strains.

## DATA AVAILABILITY STATEMENT

Short reads were deposited on SRA (accession number PRJNA1069743). Additional data (SNP data and sample annotation) are publicly available on Zenodo (10.5281/zenodo.10599894).

## AUTHOR CONTRIBUTIONS

**Marc W. Schmid:** Software, Formal analysis, Data Curation, Writing - Original draft, Writing - Review & Editing, Visualization **Aboubakr Moradi**: Conceptualization, Methodology, Investigation, Writing – Review & Editing **Deborah M. Leigh:** Conceptualization, Writing: Review & Editing **Meredith Schuman**: Methodology, Funding acquisition, Writing – Review & Editing **Sofia van Moorsel:** Conceptualization, Methodology, Investigation, Writing – Original Draft, Writing – Review & Editing, Supervision, Project administration, Funding acquisition

