## Supporting Information for "Covering the bases: population genomic structure of *Lemna minor* and the cryptic species *L. japonica* in Switzerland"

*by*

Marc W. Schmid, Aboubakr Moradi, Deborah M. Leigh,

Meredith C. Schuman, Sofia J. van Moorsel

### Table of contents

**Table S1.** Alignment rates to both reference genomes, cluster assignment, and reads per sample.

**Supplemental figure S1.** Overview field sampling.

**Supplemental figure S2.** Alignment rates for all samples

**Supplemental figure S3.** Allele frequencies in *L. minor versus L. japonica* individuals using (A) the *L. minor* reference genome (Lm7210) and (B) the *L. japonica* reference genome (Lj9421).

**Supplemental figure S4.** A) Fraction of reads assigned to bacterial, fungal, and viral sequence databases.

**Supplemental figure S5.** Comparisons of alignment rates between two reference *L. minor* clones.

**Table S1.** Alignment rates to both reference genomes, cluster assignment, and reads per sample. Delta is the difference in alignment rate between the two reference genomes.

| *Sample* | *L. minor reference genome* | *L. japonica reference genome* | *Genetic cluster* | *Delta* | *Total reads* |
|---|---|---|---|---|---|
| BA1 | 38.705 | 57.050 | B | 18.345 | 35601940 |
| BA2 | 49.980 | 74.720 | B | 24.740 | 33866577 |
| BA3 | 44.180 | 66.900 | B | 22.720 | 36067975 |
| BL2 | 55.120 | 83.520 | B | 28.400 | 33887663 |
| BL3 | 46.190 | 70.260 | B | 24.070 | 35134614 |
| BL5 | 57.665 | 86.515 | B | 28.850 | 34204498 |
| BS1 | 75.340 | 88.155 | A | 12.815 | 36929447 |
| BS2 | 74.790 | 88.663 | A | 13.873 | 34465565 |
| BS3 | 77.090 | 89.705 | A | 12.615 | 34812831 |
| CA1 | 55.245 | 78.875 | B | 23.630 | 33317664 |
| CH2 | 65.495 | 93.315 | B | 27.820 | 35261705 |
| CH3 | 65.805 | 94.140 | B | 28.335 | 36285843 |
| CH4 | 59.005 | 88.205 | B | 29.200 | 34788798 |
| CH5 | 59.047 | 86.700 | B | 27.653 | 34365591 |
| CH6 | 58.760 | 86.500 | B | 27.740 | 35359166 |
| CT1 | 49.900 | 75.185 | B | 25.285 | 35704989 |
| CT2 | 46.320 | 70.360 | B | 24.040 | 37805796 |
| CT4 | 50.080 | 73.550 | B | 23.470 | 33530103 |
| CT5 | 56.750 | 83.285 | B | 26.535 | 34561504 |
| CT6 | 56.045 | 82.605 | B | 26.560 | 34537121 |
| F4 | 53.380 | 62.130 | A | 8.750 | 39531757 |
| F5 | 46.720 | 54.000 | A | 7.280 | 33490152 |
| F6 | 48.090 | 55.540 | A | 7.450 | 40382579 |
| FR3 | 63.720 | 73.230 | A | 9.510 | 40071771 |
| GC6 | 64.835 | 93.090 | B | 28.255 | 34731668 |
| GO1 | 81.170 | 93.430 | A | 12.260 | 33658560 |
| GO2 | 73.270 | 83.950 | A | 10.680 | 35006381 |
| GO3 | 76.195 | 87.650 | A | 11.455 | 33913954 |
| GO4 | 81.645 | 94.670 | A | 13.025 | 33604080 |
| GO5 | 80.875 | 93.945 | A | 13.070 | 33171451 |
| GR3 | 59.367 | 86.540 | B | 27.173 | 35035031 |
| GR4 | 46.605 | 70.560 | B | 23.955 | 36973749 |
| GR5 | 65.270 | 92.215 | B | 26.945 | 36221215 |
| GR6 | 68.685 | 90.375 | B | 21.690 | 36553541 |
| K1 | 75.800 | 87.680 | A | 11.880 | 36412612 |
| K3_1 | 72.145 | 83.570 | A | 11.425 | 37395616 |
| K5 | 59.630 | 68.650 | A | 9.020 | 32810007 |
| KP4 | 72.635 | 82.805 | A | 10.170 | 37415990 |
| KR1 | 47.830 | 71.185 | B | 23.355 | 38173083 |
| L1 | 50.755 | 77.970 | B | 27.215 | 37216208 |
| L2 | 55.420 | 83.730 | B | 28.310 | 36671476 |
| L3 | 56.795 | 84.885 | B | 28.090 | 36799535 |
| L4 | 58.395 | 87.990 | B | 29.595 | 37377588 |
| L5 | 58.680 | 87.750 | B | 29.070 | 39744236 |
| L6 | 59.320 | 88.870 | B | 29.550 | 36998414 |
| LB1 | 30.540 | 46.875 | B | 16.335 | 37063681 |
| LB2 | 38.605 | 58.935 | B | 20.330 | 34493492 |

| | | | | | |
|---|---|---|---|---|---|
| LB4 | 33.150 | 51.450 | B | 18.300 | 40846825 |
| LB5 | 44.040 | 66.570 | B | 22.530 | 32729501 |
| Lemna_9478 | 25.755 | 91.900 | outgroup | 66.145 | 37420333 |
| Lemna_9700 | 2.650 | 2.910 | outgroup | 0.260 | 33610340 |
| Lemna_9965 | 59.770 | 91.425 | B | 31.655 | 37860161 |
| Lemna_9967_2 | 82.790 | 95.700 | A | 12.910 | 38115442 |
| Lemna_9969_1 | 62.935 | 95.355 | B | 32.420 | 34811870 |
| Lemna_9978_2 | 61.080 | 93.520 | B | 32.440 | 31879455 |
| Lemna_9983 | 63.580 | 95.500 | B | 31.920 | 33172894 |
| LR1 | 71.050 | 81.720 | A | 10.670 | 34689338 |
| LR5 | 81.847 | 92.143 | A | 10.297 | 35576834 |
| LR6 | 78.620 | 89.450 | A | 10.830 | 35763846 |
| M2 | 61.330 | 70.435 | A | 9.105 | 34116778 |
| M3 | 64.510 | 74.140 | A | 9.630 | 35224815 |
| M4 | 64.020 | 73.980 | A | 9.960 | 48889332 |
| M5 | 67.675 | 77.590 | A | 9.915 | 34515526 |
| M6_1 | 60.327 | 69.130 | A | 8.803 | 33627159 |
| M6_2 | 60.210 | 69.070 | A | 8.860 | 33892364 |
| MT1 | 57.250 | 85.780 | B | 28.530 | 34547409 |
| MT2 | 57.105 | 85.785 | B | 28.680 | 34229283 |
| MT3 | 62.595 | 92.295 | B | 29.700 | 35699159 |
| MT4 | 58.365 | 87.945 | B | 29.580 | 33948058 |
| MT6 | 57.520 | 89.390 | B | 31.870 | 36846654 |
| R4 | 61.410 | 70.750 | A | 9.340 | 57006716 |
| R5 | 33.320 | 50.880 | B | 17.560 | 38553680 |
| R6 | 33.060 | 50.160 | B | 17.100 | 32674827 |
| SD1 | 59.340 | 85.720 | B | 26.380 | 34699347 |
| SD2 | 60.915 | 89.185 | B | 28.270 | 36359607 |
| SD3 | 53.890 | 78.040 | B | 24.150 | 33340968 |
| SD4 | 57.550 | 83.500 | B | 25.950 | 37362791 |
| SD5 | 53.850 | 77.123 | B | 23.273 | 37309360 |
| SD6 | 57.870 | 82.820 | B | 24.950 | 33619084 |
| TO1 | 54.470 | 62.590 | A | 8.120 | 34509384 |
| TO2 | 40.937 | 46.577 | A | 5.640 | 38256580 |
| Y2_1 | 50.060 | 57.830 | A | 7.770 | 38650611 |
| Y3 | 73.560 | 85.180 | A | 11.620 | 37005581 |
| Y4 | 65.270 | 75.385 | A | 10.115 | 35922842 |
| Y5 | 61.460 | 70.765 | A | 9.305 | 34833359 |
| Y6 | 60.040 | 68.880 | A | 8.840 | 32999359 |
| Z3 | 67.425 | 78.350 | A | 10.925 | 34928266 |
| Z4 | 71.400 | 83.100 | A | 11.700 | 46203913 |
| Z5 | 53.675 | 61.740 | A | 8.065 | 38994208 |

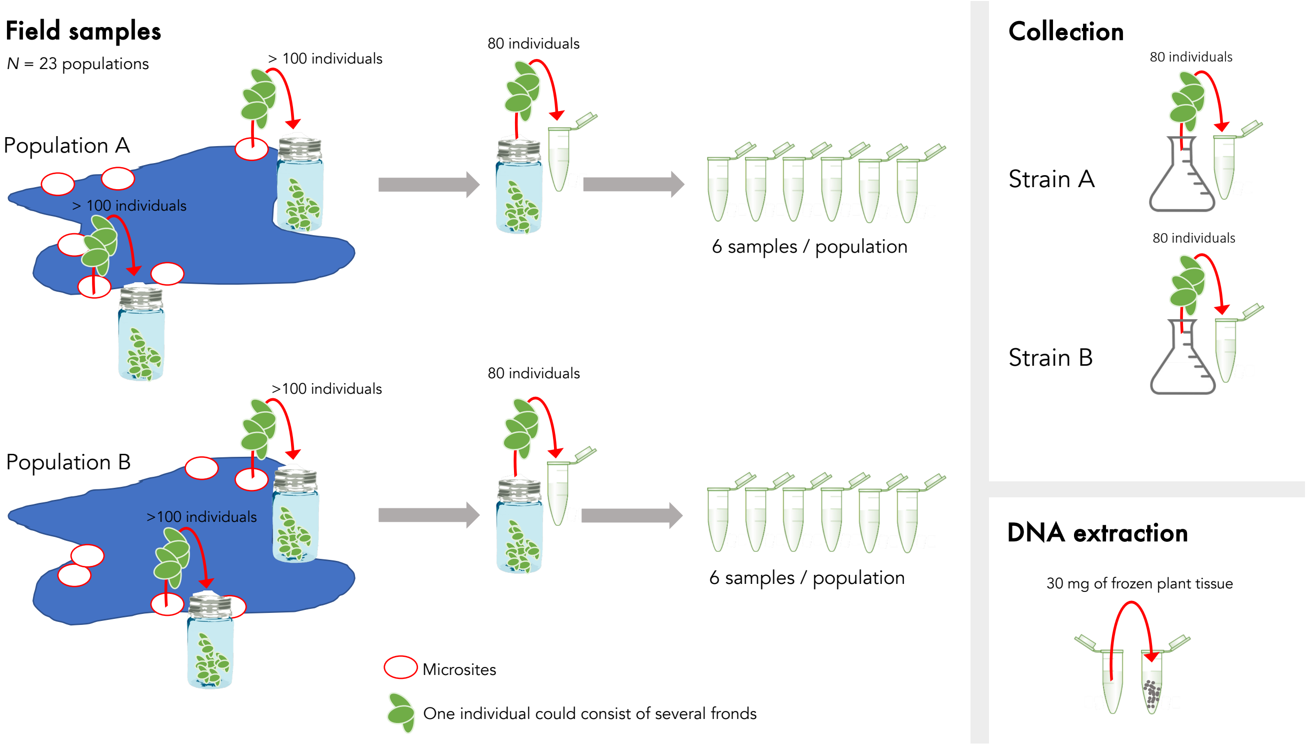


**Supplemental figure S1.** Overview field sampling.

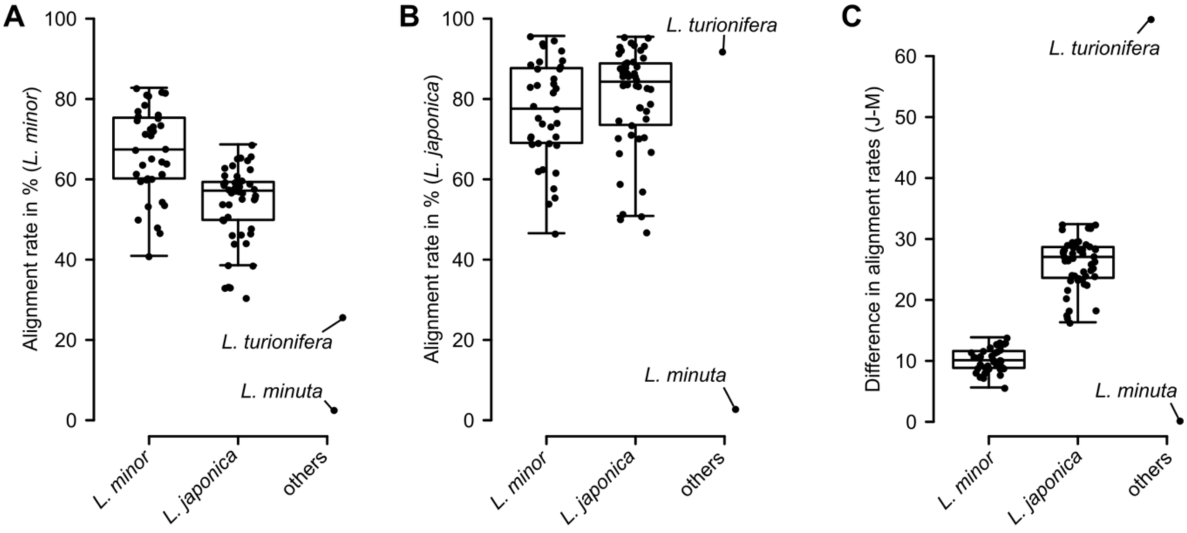


**Supplemental figure S2.** Alignment rates for all samples using the *L. minor* (A) reference genome (strain 7210) and the *L. japonica* (B) reference genome (strain 9421).C) Difference between (B) and (A) (*L. japonica* - *L. minor*). The groups (*L. minor* and *L. japonica*) were inferred from the kinship matrices in Figure 2A.

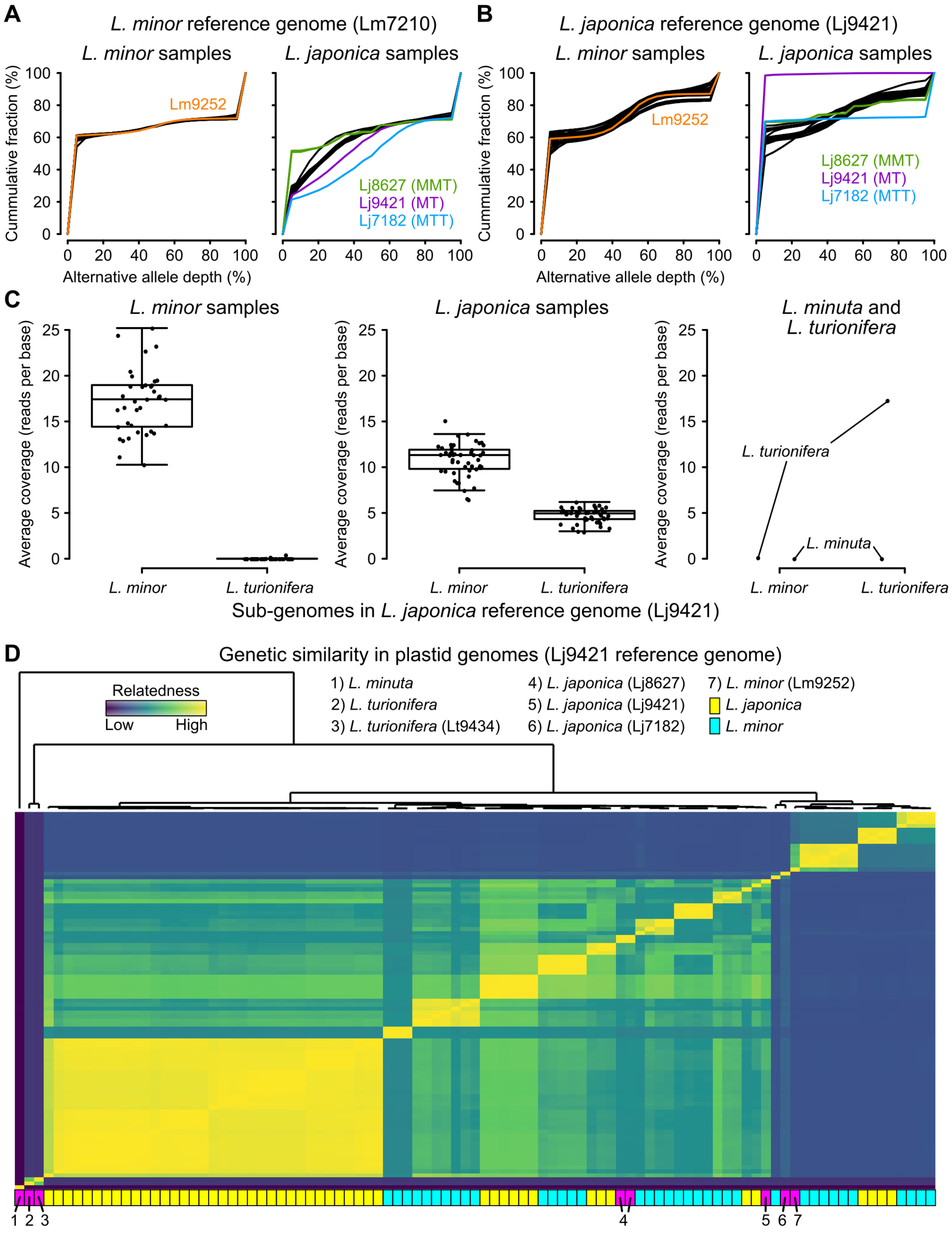
A
L. minor reference genome (Lm7210)
L. minor samples
L. japonica samples
Cummulative fraction (%)
Lm9252
Lj8627 (MMT)
Lj9421 (MT)
Lj7182 (MTT)
Alternative allele depth (%)
B
L. japonica reference genome (Lj9421)
L. minor samples
L. japonica samples
Cummulative fraction (%)
Lm9252
Lj8627 (MMT)
Lj9421 (MT)
Lj7182 (MTT)
Alternative allele depth (%)
C
L. minor samples
L. japonica samples
L. minuta and L. turionifera
Average coverage (reads per base)
L. turionifera
L. minuta
L. minor
L. turionifera
Sub-genomes in L. japonica reference genome (Lj9421)
D
Genetic similarity in plastid genomes (Lj9421 reference genome)
Relatedness
Low
High
1) L. minuta
2) L. turionifera
3) L. turionifera (Lt9434)
4) L. japonica (Lj8627)
5) L. japonica (Lj9421)
6) L. japonica (Lj7182)
7) L. minor (Lm9252)
L. japonica
L. minor
1 2 3
4
5 6 7

**Supplemental figure S3.** Allele frequencies in *L. minor versus L. japonica* individuals using (A) the *L. minor* reference genome (Lm7210) and (B) the *L. japonica* reference genome (Lj9421). Highlighted samples were previously published data: Lm9252 was a diploid *L. minor* individual (i.e., not double haploid as the individual that was used for the reference genome, Lm7210). The allele frequencies found in this sample were similar to the ones observed in our *L. minor* samples (black lines). Lj9421, Lj7182, and Lj8627 were three different *L. japonica* individuals with different genome ratios. Lj9421 used for the reference genome was diploid with one copy of each, the *L. minor* and *L. turionifera*, genomes. Lj7182 and Lj8627 were triploid with either one or two copies of the *L. minor* genome, respectively. The lower the *L. minor* content, the higher the alternative allele ratios. Our samples matched best to Lj8627 with only one *L. minor* genome copy and two copies of the *L. turionifera* genomes. This was clearer when using the *L. japonica* reference genome. C) Average read coverage of the *L. minor* and *L. turionifera* sub-genomes of the *L. japonica* reference genome Lj9421. D) Visualization of genetic similarities in the plastid genomes using all samples including publicly available data. The latter and the outgroup species *L. minuta* and *L. turionifera* are highlighted in magenta at the bottom.

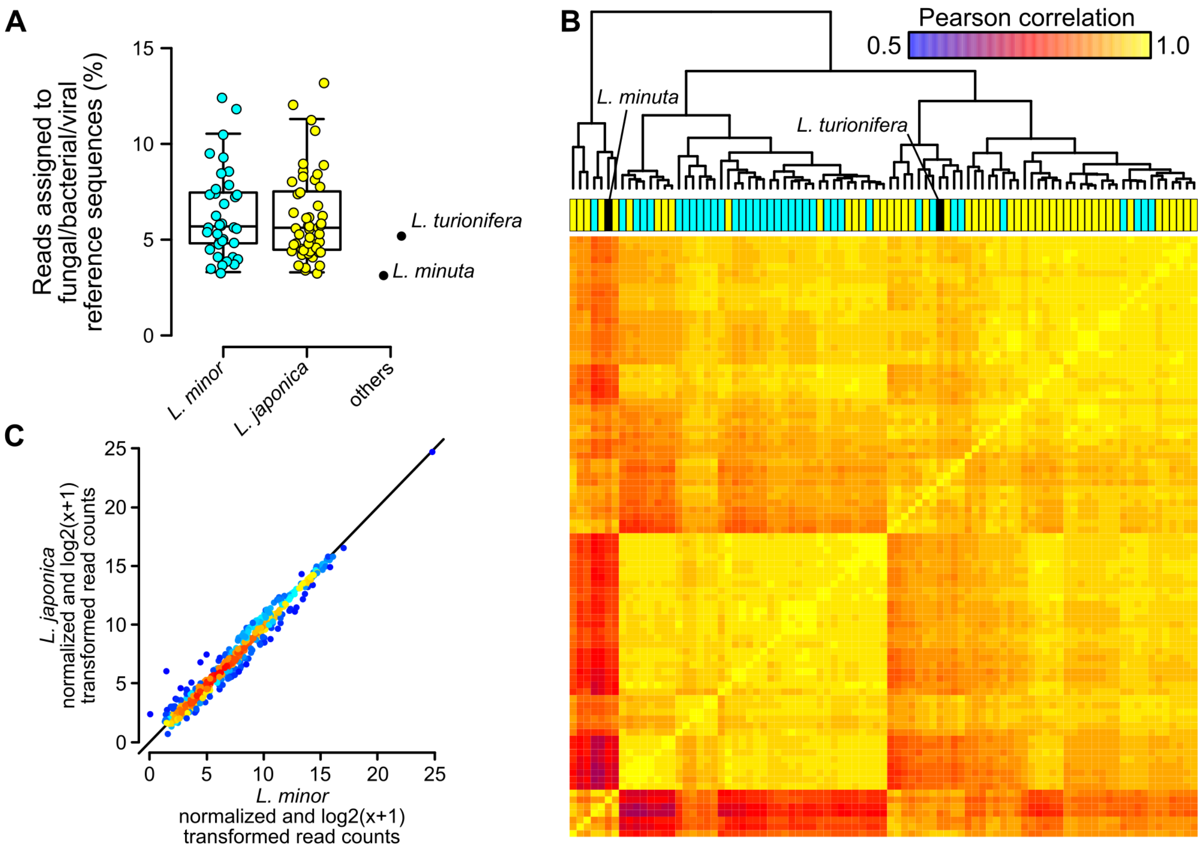


**Supplemental figure S4.** A) Fraction of reads assigned to bacterial, fungal, and viral sequence databases. B) Pearson correlation matrix and dendrogram using the log2(x+1)-transformed and normalized read counts per bacterial/fungal/viral family (n = 555). C) Scatterplot with the average *L. minor* and *L. japonica* samples (n = 555).

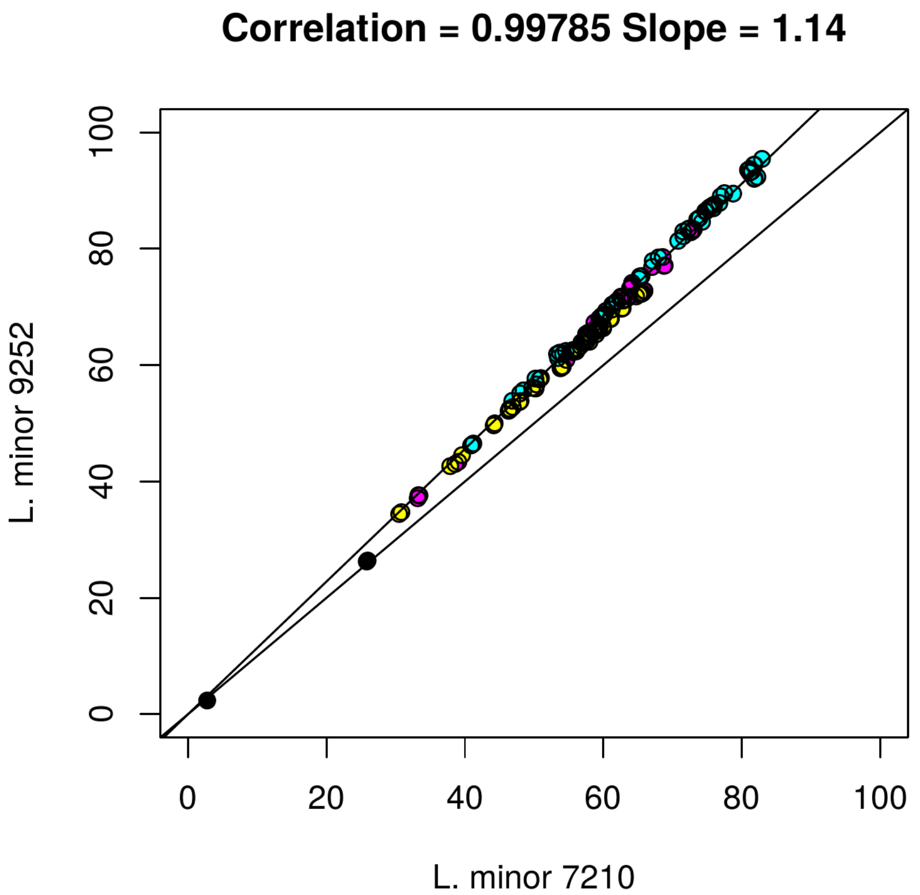


**Supplemental figure S5.** Comparisons of alignment rates between two reference *L. minor* clones. The phylogenetically and geographically closer reference *L. minor* 9252 (Haltiala, Finland), rather than *L. minor* 7210 (Makhanda, S. Africa) shows slightly improved alignment rates but conclusions are not affected. Alignment rates with 9252 were about 14 % higher than with 7210, however it affected all samples linearly. The Pearson correlation is 0.998.
